# Characterization of the Novel Mitochondrial Genome Segregation Factor TAP110 in *Trypanosoma brucei*

**DOI:** 10.1101/2020.06.25.171090

**Authors:** Simona Amodeo, Ana Kalichava, Albert Fradera-Sola, Eloïse Bertiaux-Lequoy, Paul Guichard, Falk Butter, Torsten Ochsenreiter

## Abstract

Proper mitochondrial genome inheritance is key for eukaryotic cell survival, however little is known about the molecular mechanism controlling this process. *Trypanosoma brucei*, a protozoan parasite, contains a singular mitochondrial genome aka kinetoplast DNA (kDNA). kDNA segregation requires anchoring of the genome to the basal body via the tripartite attachment complex (TAC). Several components of the TAC as well as their assembly have been described, it however remains elusive how the TAC connects to the kDNA. Here, we characterize the TAC associated protein TAP110 and for the first time use ultrastructure expansion microscopy in trypanosomes to reveal that TAP110 is the currently most proximal kDNA segregation factor. The kDNA proximal positioning is also supported by RNAi depletion of TAC102, which leads to loss of TAP110 at the TAC. Overexpression of TAP110 leads to expression level changes of several mitochondrial proteins and a delay in the separation of the replicated kDNA networks. In contrast to other kDNA segregation factors TAP110 remains only partially attached to the flagellum after DNAse and detergent treatment and can only be solubilized in dyskinetoplastic cells, suggesting that interaction with the kDNA might be important for stability of the TAC association. Furthermore, we demonstrate that the TAC, but not the kDNA, is required for correct TAP110 localization *in vivo* and suggest that TAP110 might interact with other proteins to form a >669 kDa complex.

**Summary Statement:** TAP110 is a novel mitochondrial genome segregation factor in *Trypanosoma brucei* that associates with the previously described TAC component TAC102. Ultrastructure expansion microscopy reveals its proximal position to the kDNA.

## Introduction

Mitochondria are a defining feature of eukaryotic cells. They perform a large number of different functions ranging from catabolic reactions like oxidative phosphorylation [1] to anabolic processes like iron sulfur cluster assembly [2] and calcium homeostasis [3]. The vast majority of the mitochondrial proteins are encoded and expressed from the nuclear genome, while only a small set of proteins, mostly of the oxidative phosphorylation chain are encoded on the organelle’s own genome. . In *Trypanosoma brucei* a parasitic protist the mitochondrial genome is organized in a complex structure named kinetoplast DNA (kDNA). It consists of about 25 large (23 kbp) circular DNA molecules [4] that encode 16 genes of the oxidative phosphorylation chain, two ribosomal proteins [5] and two ribosomal RNAs. Twelve of the mitochondrial genes require posttranscriptional modifications by RNA editing prior to translation on the mitochondrial ribosomes [6–9]. The guide RNAs involved in this process are encoded on minicircles (1 kbp) of which about 5000 are catenated into the kDNA network forming a disc like structure [10]. In that network the minicircles are oriented perpendicular to the horizontal plane of the disc [11–14]. The maxicircles are interwoven into the minicircle network and also interlocked with each other [4,15]. Replication of the kDNA occurs during G1 of the parasite cell cycle, just prior to start of nuclear DNA replication. Our current model of kDNA replication predicts that the minicircles are released by a not identified topoisomerase activity from the network into the kinetoflagellar zone (KFZ), the region between the inner mitochondrial membrane and the kDNA disc [16] [17]. Here the replication is initiated through binding of the Universal Minicircle Sequence Binding Protein (UMSBP) [18], a primase activity [19] and a bacterial polI like enzyme (POLIB, [20,21]). Replication proceeds unidirectional via theta intermediates [22]. The replication products are subsequently separated and transported by an unknown mechanism to the opposing ends of the kDNA disc. At the antipodal sites the RNA primers are removed, most of the minicircle gaps are repaired and the replicated DNA is reattached to the growing kDNA disc [17] [23]. Maxicircle replication is much less well understood but likely also proceeds via theta intermediates and occurs while the maxicircles are attached to the kDNA disc [24]. Once all minicircles have been replicated and reattached the remaining gaps in the network are sealed [25,26] and the two daughter networks are segregated through the movement of the basal bodies of the flagellum [27]. The physical connection between the kDNA and the basal bodies that mediates the segregation was described in electron microscopy studies and was termed the tripartite attachment complex (TAC). The TAC consists of (i) the exclusion zone filaments, a region between the basal bodies and the outer mitochondrial membrane that is devoid of ribosomes, (ii) the differentiated mitochondrial membranes and (iii) the unilateral filaments that connect the inner mitochondrial membrane to the kDNA [28]. Several proteins of this structure have been characterized [23,29]. TAC102 is the kDNA most proximal TAC component known. however it remains unclear if TAC102 binds directly to kDNA disc or if other proteins are mediating this process [30,31]. The closest interactor of TAC102 is the transmembrane domain containing protein p166 that is localized at the inner mitochondrial membrane [32]. Three outer mitochondrial membrane components of the TAC (TAC40, TAC42 and TAC60, [33,34]) as well as two components in the exclusion zone filaments (p197, TAC65, [34–36]) are essential for proper kDNA segregation. While TAC65 is localized close to the outer mitochondrial membrane the second component of the EZF (P197) is close to the basal bodies of the flagellum. Additionally, there are a number of proteins that beside their role in the TAC have other functions like TbTBCCD1, pATOM36, alpha-KDE2 and AEP1 [37–39]. The overall complex is assembled *de novo* in a hierarchical manner from the maturing basal body towards the kDNA during G1 of the trypanosome cell cycle [36]. While we and others have identified components of all three TAC regions, it remains unknown how and through which components the TAC is connected to the kDNA. In order to identify novel components of the TAC that might interact with the kDNA we used an N-terminally tagged TAC102 as bait to purify interacting partners. Here we present the results of a novel protein TAP110 that based on ultrastructure expansion microscopy (U-ExM) is proximal to the kDNA relative to TAC102.

## Results

TAP110 (Tb927.11.7590) is a basic (pI = 8), 110 kDa, hypothetical conserved protein with a predicted mitochondrial targeting sequence at the N-terminus (Figure S1A). The C-terminal part of TAP110 shows similarities to the micronuclear linker histone polyprotein of *Tetrahymena thermophila*. Furthermore, TAP110 contains six posttranslational modifications in form of methylated arginines (Figure S1A) [40]. TAP110 has been detected in TAC102 immunoprecipitation analyses, where N-terminal tagged versions of TAC102 were used to co-immunoprecipitate potential interaction partners (Figure S1B+C). The most abundant protein in the eluate was TAC102, the second and third most enriched proteins were Tb927.11.6660 and TAP110. Based on a phylogenetic comparison TAP110 shares a common evolutionary history with TAC102 (Figure S1D, also see discussion).

### TAP110 localization

To localize the TAP110 protein, we tagged it *in situ* at the C-terminus with a PTP epitope tag in New York single marker (NYsm) bloodstream form (BSF) *T. brucei [41,42]*. We performed immunofluorescence microscopy and used an anti-Protein A antibody to detect TAP110-PTP. Based on colocalization studies with the basal body marker YL1/2 and the DNA stain DAPI, the protein localizes between the kDNA and the basal bodies (Figure 1A). TAP110 localization during the cell cycle resembled the typical localization pattern of a TAC component, where two signals for TAP110 are discernable before the kDNA is segregated but only after the separation and maturation of the daughter basal body. (arrowheads Figure 1A). Based on the proximity to TAC102 and the kDNA in epifluorescence microscopy we decided to compare TAP110 and TAC102 localization by super resolution microscopy using stimulated emission depletion (STED) microscopy. Based on 16 images with a minimum X-Y resolution of 37.9 nm, TAP110 and TAC102 co-localize with a Pearson correlation coefficient of 0.841 (n = 16) (Figure 1B).

**Figure 1:**
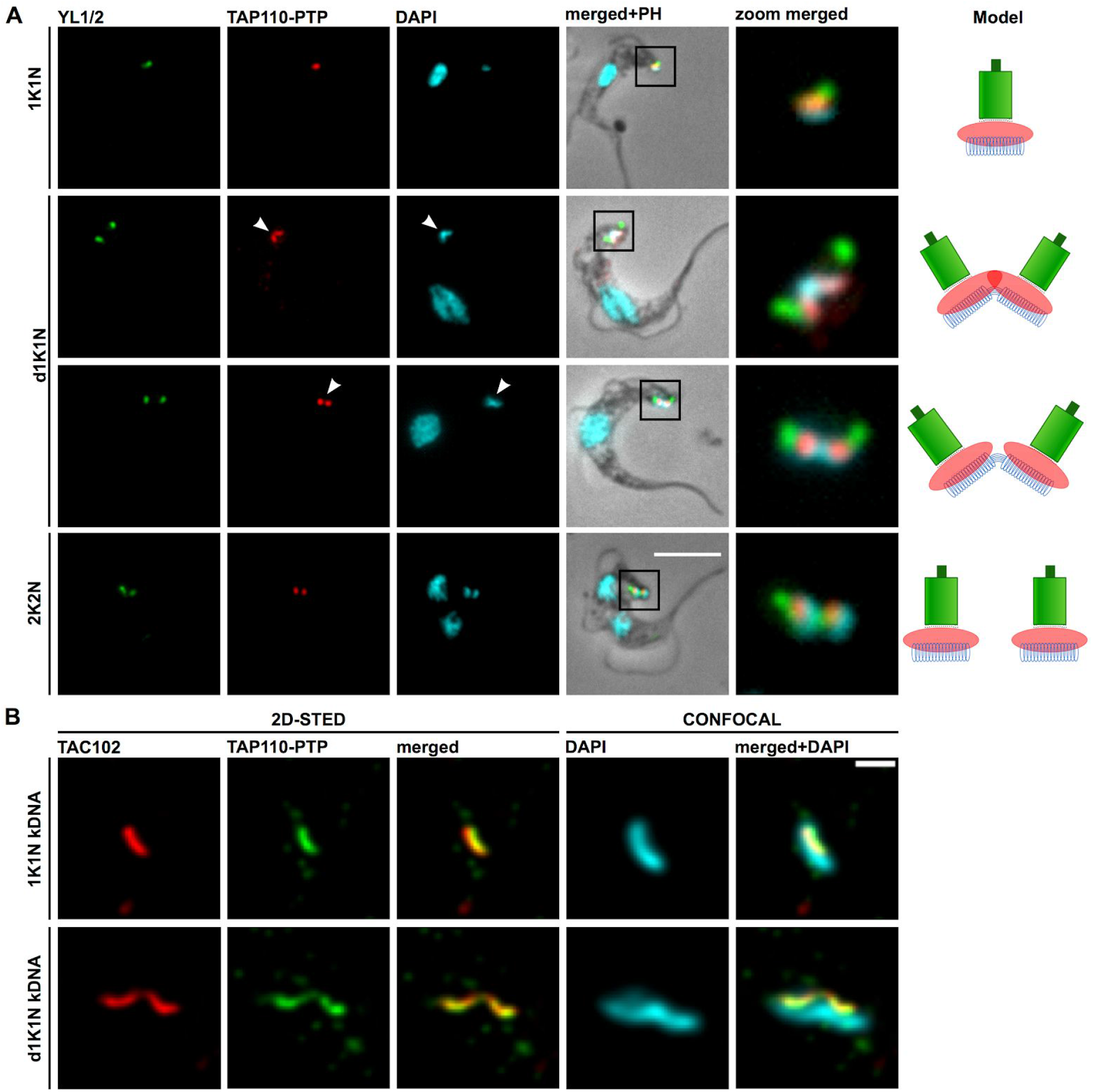
Localization of TAP110 in *T. brucei* BSF cells. **A)** Representative immunofluorescence microscopy images of TAP110-PTP expressing BSF cells during different stages of the cell cycle (1K1N, dK1N, 2K2N). The mature basal bodies (green) were detected with the YL1/2 monoclonal antibody. TAP110-PTP (red) was detected by means of the anti-Protein A antibody. The kDNA and the nucleus were stained with DAPI (cyan). The panel on the right side shows a simplified model of TAP110 localization during the cell cycle. Green depicts the basal bodies, red TAP110 and blue the kDNA. Scale bar: 5 μm. Zoom factor: 4x. Arrowheads point towards duplicated TAP110 signals and duplicated kDNAs. **B)** Deconvoluted 2D-STED immunofluorescence images of TAP110-PTP and TAC102 expressed in BSF cells. TAC102 (red) was detected with the anti-TAC102 monoclonal antibody, TAP110-PTP (green) and the kDNA (cyan) were detected as described above. The TAC102 and TAP110 signal were acquired by 2D-STED, the kDNA by confocal microscopy. Scale bar: 500 nm; dK, duplicating kDNA; K, kDNA; N, nucleus; PH, phase contrast.

### Depletion of TAP110 by RNAi

To study the function of TAP110 we depleted the mRNA by RNAi in NYsm BSF cells that contained an endogenously tagged allele of TAP110 (as described above) using a tetracycline (tet) inducible RNAi vector [43]. The knock-down efficiency of TAP110 mRNA was monitored by probing for the PTP-tagged TAP110 protein in western blots from whole cell protein extracts. Depletion of TAP110 was efficient but not complete (Figure 2A). Loss of TAP110 did not lead to a detectable growth defect or morphological changes of the cells (Figure 2B). In order to test if TAP110 depletion has an effect on the TAC we probed the western blot for the potential interactor TAC102 but didn’t observe a loss of the signal (Figure 2A).

**Figure 2:**
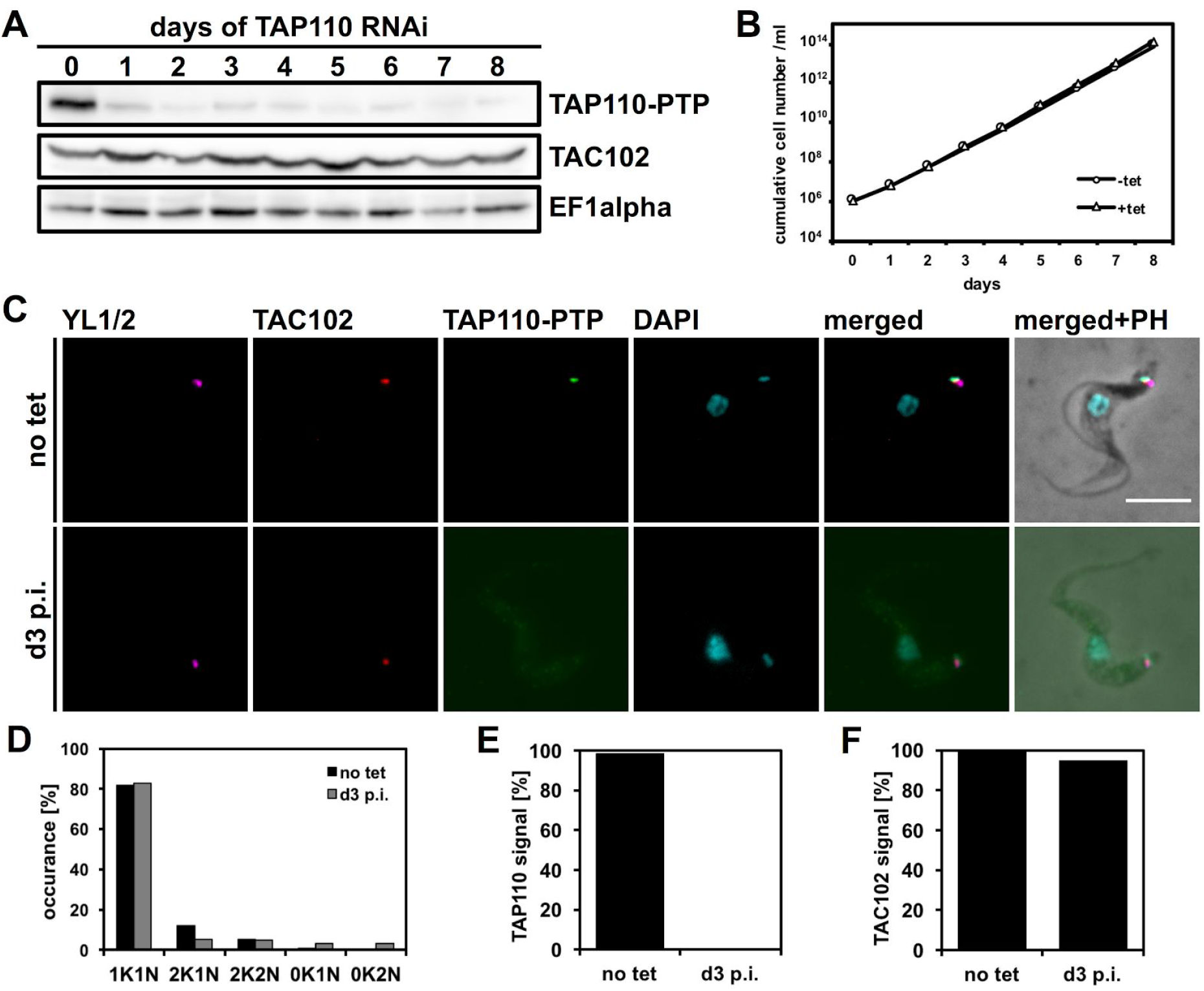
Phenotypes upon depletion of TAP110 mRNA by RNAi in BSF cells. **A)** Western blot showing depletion of TAP110–PTP protein at different days of the RNAi induction. TAP110-PTP was detected by rabbit IgGs and TAC102 was detected using a monoclonal anti TAC102 antibody. EF1alpha serves as a loading control. **B)** Growth curve of tet inducible BSF TAP110 RNAi TAP110-PTP cells. **C)** Immunofluorescence images of non-induced cells (no tet) and cells at day three post induction (d3 p.i.). The signals are represented by maximum intensity projections from image stacks. Basal bodies, TAP110-PTP, TAC102 and DNA were detected as described in Figure 1. Scale bar: 5 μm **D)** Quantification of the relative occurrence of kDNA networks and nuclei of non-induced cells (no tet) and cells at day three post-induction (d3 p.i.). N≥100 cells for each condition were analyzed. **E)** Quantification of TAP110 signal from the experiment shown in C). **F)** Quantification of TAC102 signal as performed in E). K, kDNA; N, nucleus; PH, phase contrast.

In order to characterize a potential effect of TAP110 depletion on the mitochondrial genome and TAC102 we performed immunofluorescence microscopy with TAP110 depleted BSF cells at day three of RNAi (Figure 2C). The basal body, TAP110 and the DNA were detected as described in Figure 1. TAC102 was detected with a monoclonal anti-TAC102 antibody. We sampled the population (n≥100 for each condition) at day zero (no tet) and three days post induction (d3 p.i.) and evaluated the relative occurrence of kDNA networks and nuclei in different cell cycle stages: cells with one kDNA network and one nucleus (1K1N; cells are in G1 of the cell cycle), cells with already replicated and segregated kDNA networks and one nucleus (2K1N, cells are in nuclear G2 phase), as well as cells that had replicated and segregated both the kDNA and the nucleus (2K2N, cells just prior to cytokinesis). We also screened for abnormal K-N combinations like 0K1N as well as 0K2N (indicative of kDNA replication/segregation defects). We did not observe a change in K-N combinations between non-induced and induced cells at day three of the RNAi (Figure 2D). We also analysed the effect of TAP110 RNAi on the kDNA at day six and found the networks to be 15% increased in DNA content as measures by DAPI staining ((n>200 cells, p ≤ 0.001) Figure S2). We further quantified the TAP110 signal, which was not detectable at the third day of TAP110 RNAi (Figure 2E). For TAC102 we observed a loss of signal in around 5% of the induced cells (Figure 2F). Furthermore, 8% of the induced cells showed a weaker signal for TAC102.

### Overexpression of TAP110

To further evaluate the function of TAP110 we induced overexpression of a triple HA tagged ectopic version of TAP110 in PCF cells (29-13) and monitored cell growth and abundance of TAP110-HA (Figure 3A+B). Furthermore, we used immunofluorescence microscopy to analyse effects of TAP110-HA overexpression on the TAC and the mitochondrial genome (Figure 3C). As a proxy for TAC presence we used an anti-TAC102 antibody, while the kDNA was detected with DAPI. We observed an accumulation of cells with replicated, non-segregated kDNA networks (dK1N), which was most dominant at day two of the overexpression, where almost 50% of the cells had a duplicated non-segregated kDNA (Figure 3D), while this was about 20% in the wild type cells.

**Figure 3:**
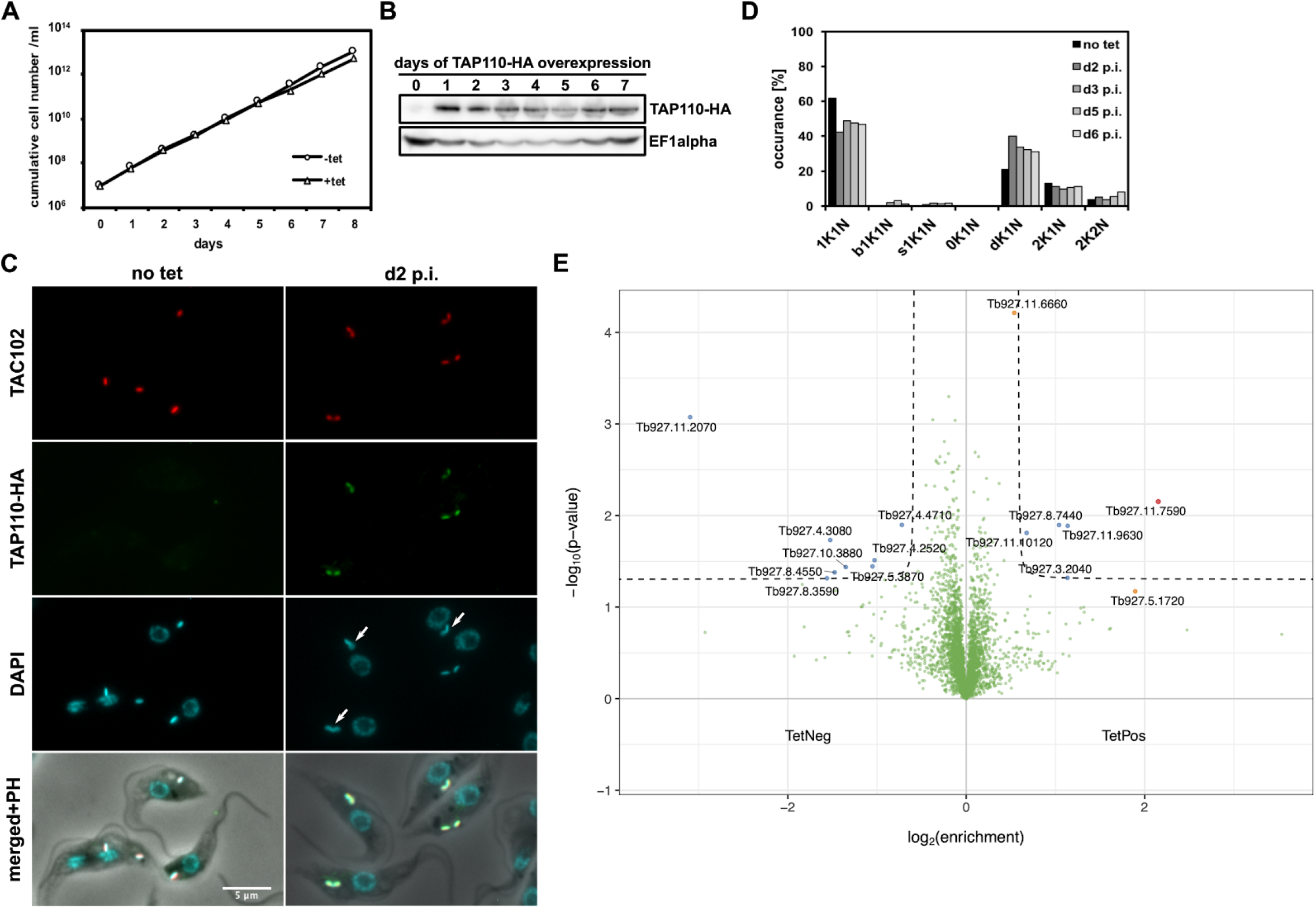
Phenotypes upon overexpression of TAP110-HA in PCF cells. **A)** Growth curve of tet inducible PCF TAP110-HA cells. **B)** Western blot showing expression of TAP110-HA protein at different days of overexpression. TAP110-HA was detected by an anti-HA antibody; EF1-alpha serves as a loading control. **C)** Immunofluorescence microscopy images of non-induced (no tet) and induced cells at day two post induction of the overespression construct (d2 p.i.). TAC102 and DNA were detected as described in Figure 1 and TAP110-HA was detected by an anti-HA antibody. Scale bar: 5 μm. Arrows point towards duplicated non-segregated kDNAs **D)** Quantification of the relative occurrence of kDNA networks and nuclei before inducing the overexpression (no tet) and at different days post induction (d2-d6 p.i.). N≥100 cells for each condition were analyzed. **E)** Volcano plot of proteins in tet positive (d2 p.i.) against tet negative cells. Highlighted in red is TAP110 (enrichment 4.45), the proteins highlighted in blue are possible interactors passing the threshold of a p-value <0.05 and log2FoldChange >1 or <−1. Further highlighted in orange are possible interactors not passing the threshold of p-value <0.05 or log2FoldChange >1 or <−1. bK, big kDNA; dK, duplicating kDNA; K, kDNA; N, nucleus; PH, phase contrast; sK, small kDNA.

We also analysed the overall proteome changes using mass spectrometry in cells where TAP110 was overexpressed. For this we induced expression of TAP110-HA for two days and then harvested whole cells and compared the total cell proteome to cells not induced for TAP110-HA overexpression (Figure 3E). TAP110, highlighted in red, was found 4.45 fold enriched after induction with textracyclin (d2 p.i.). Twelve proteins were differentially expressed during the TAP110 overexpression (highlighted in blue) and passed the threshold of a p-value <0.05 and a fold change −1 < log_2_> 1. Of these, eight were depleted and four were upregulated. Further we identified two proteins upregulated passing one of the thresholds (either log_2_ > 1 or p-value < 0.05) and being close to pass the other (highlighted in orange).

Of the proteins depleted during TAP110 overexpression, four were annotated as conserved hypothetical proteins. Tb927.11.2070 is likely an essential protein and has an IEP of 7.96. Tb927.8.3590 has an IEP of 9.95, both are not predicted to contain a conventional mitochondrial targeting signal [44]. Tb927.4.4710 and Tb927.10.3880 had more neutral or acidic IEPs, had as well no conventional mitochondrial targeting signal predicted and were both annotated as non-essential, too. The other four depleted proteins were all likely to be mitochondrial. Tb927.4.3080 is a basic (PI 8.45) putative uncharacterized ADAM cysteine-rich domain YggU family protein that is likely imported into the mitochondrion [45]. Then Tb927.8.4550 and Tb927.5.3870 are both putative mitochondrial ribosomal proteins of the small and the large subunits, respectively. Tb927.4.2520 is the silent information regulator 2 related protein 3, shown to be localized at the kDNA [46,47].

Of the six proteins increased in abundance, one (Tb927.11.10120) is a conserved hypothetical protein with an IEP of 10.27 but likely not a mitochondrial protein [44]. Tb927.3.2040 is a putative kinesin motor protein localized to the hook complex. Tb927.11.9630 is a putative vacuolar-sorting protein. N-terminal tagging shows that it has an endocytic localization pattern [47]. Tb927.8.7440 is a putative lipase/hydrolase showing a similar endocytic localization pattern as Tb927.11.9630. The other two upregulated proteins passing one of the two thresholds, Tb927.5.1720 and Tb927.11.6660 (Figure S5) are both associated with the kDNA [47]. Tb927.11.6660 was initially identified as an interactor of TAC102 (Figure S1). While Tb927.11.6660 is present at the kDNA throughout the cell cycle, it was observed in the nucleus only after start of nuclear S phase (Figure S5).

Since the cell growth was not affected by the depletion (Figure 2) and overexpression of TAP110 only had a minor effect on cell growth (Figure 3), we wanted to test the fitness of TAP110 depleted and overexpressing cells during stress condition. For this, we decided to apply commonly used stress conditions i.e heat stress (33°C) for PCF cells and EtBr treatment for BSF cells. In the EtBr stress condition non-induced and and induced cells performed the same (Figure S3A-D). In the heat stress condition on the other hand, we observed a slightly stronger growth retardation in induced cells grown at 33°C (compare Figure 3A and S3E). When we analysed DAPI stained cells at day eight post induction, we observed an increased number of cells with abnormal kDNA network content (Figure S3F).

### Impact of of TAC102 depletion on TAP110

To further analyse potential interactions of TAP110 with the TAC, we investigated the effect of TAC102 RNAi on TAP110. Previous studies have shown that the TAC is assembled hierarchically from the base of the flagellum to the kDNA [36]. Consequently, depletion of a TAC protein distal to the kDNA leads to loss of the kDNA proximal TAC components [36]. Thus, if TAP110 is closer to the kDNA than TAC102, depletion of the latter should lead to a loss of TAP110. We performed immunofluorescence microscopy on cells from day three post TAC102 depletion (d3 p.i.) (Figure 4A). In addition to the depletion of the TAC102 signal and the previously described kDNA loss/segregation phenotype [30] we also observed that more than 80% of the induced cells had lost the signal for TAP110 (Figure 4B-C). We also probed for TAC102 and TAP110 in a western blot (n = 3) and found that both protein where significantly depleted (TAC102 <10% remaining; TAP110 <50% remaining; (Figure 4D-E)).

**Figure 4:**
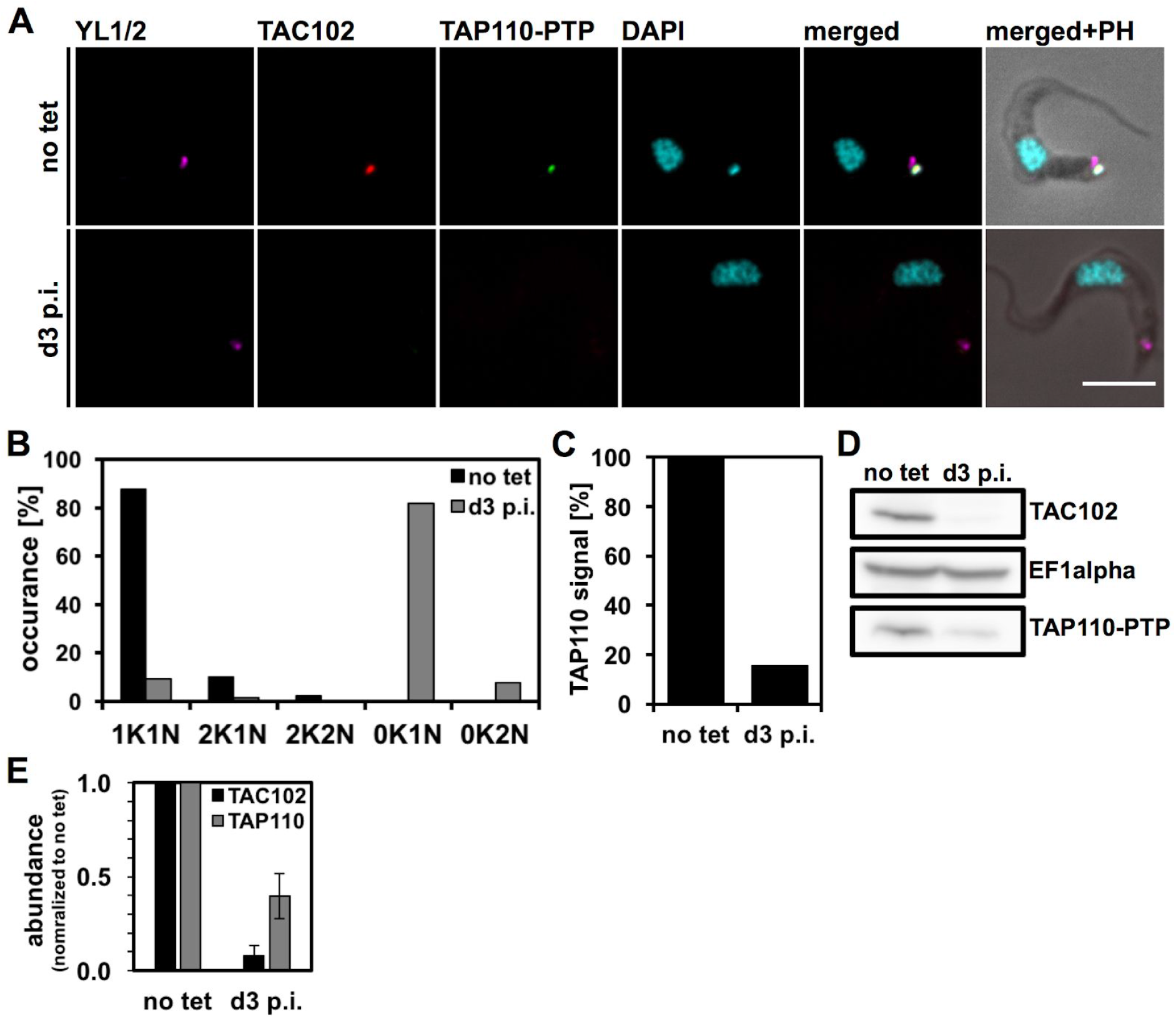
Effect of TAC102 depletion on TAP110 localization and abundance. **A)** Immunofluorescence microscopy images of non-induced (no tet) and induced cells at day three post-induction (d3 p.i.). Images were produced as described in Figure 1. Scale bar: 5 μm. **B)** Quantification of the relative occurrence of kDNA networks and nuclei before inducing the RNAi (no tet) and at day three post induction (d3 p.i.). A Minimum of n≥100 cells for each condition were analyzed. **C)** Quantification of TAP110 signal from the experiment shown in A). **D)** Western blot showing depletion of TAC102 and TAP110-PTP at day three of RNAi (d3 p.i.). Probing for EF1alpha serves as a loading control. **E)** Quantification of TAP110 and TAC102 signals from blot seen in D). K, kDNA; N, nucleus; PH, phase contrast.

### Ultrastructure expansion microscopy

Based on the TAC102 RNAi experiments we predicted TAP110 to be proximal to the kDNA, however this was not confirmed by superresolution microscopy (see Figure 1B). In order to further improve the resolution we established ultrastructure expansion microscopy for insect form *T. brucei* cells.

Immunostaining with an anti-tubulin antibody in combination with confocal microscopy showed that the PCF trypanosome cells were “equally” expanded in all three dimensions, largely retaining the trypomastigote morphology of an elongated cell body that tapers at the anterior and posterior end (Figure 5A, Figure S4). To further investigate the expansion process inside the cell, we stained nucleus and kDNA with DAPI and compared non-expanded with expanded cells. We observed isotropic expansion of the nucleus by a factor of 3.86 +/− 0.594 (n=22, Figure 5B-D), while the expansion of the kDNA disc was largely limited to the planar axis of the disc, increasing the diameter by a factor of 3.75 +/− 0.628 (Figure 5B-D). As a third parameter, we measured the expansion of the basal body diameter and compared it to thin section electronmicroscopy imagery (n=12) from non-expanded chemically fixed cells. We found the basal body to be isotropically expanded by a factor of 3.61 +/− 0.14 (Figure 5B-D, Figure S4).

**Figure 5:**
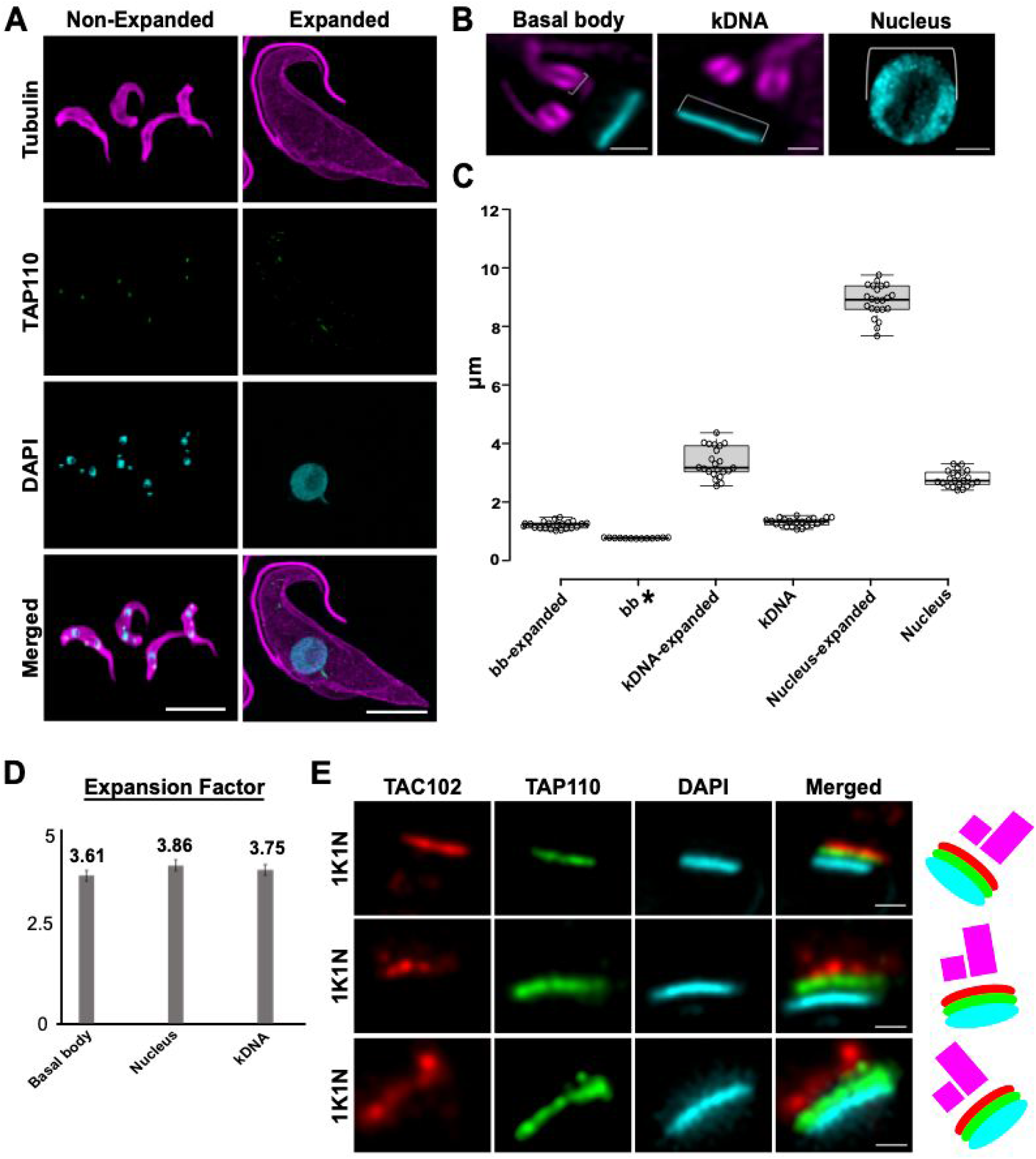
*T. brucei* expansion with U-ExM. **A)** Non-expanded and expanded PCF cells stained with α-tubulin (magenta; Alexa Fluor 594), TAP110 (green; Oregon Green 488), kDNA (cayan; DAPI) and imaged by confocal microscopy followed by deconvolution. Scale bar: 20 μm. **B)** Magnified views of the basal body, kDNA and nucleus. Scale bar: 2.5 μm. **C)** Measurements of basal body size, kDNA length and nucleus diameter in non-expanded and expanded cells (expanded basal bodies, kDNAs and nuclei each n=22 cells; non-expanded kDNAs and nuclei each n=22 cells; * corresponds to non-expanded basal body measurements from TEM (n=12 cells)). **D)** Expansion factor calculated as the ratio between non-expanded and expanded basal body, kDNA and nucleus from measurements obtained in C. **E)** Representative images of localized TAP110 (green; Oregon Green 488), TAC102 (red, Alexa Fluor 594) and kDNA (cayan, DAPI) in expanded cells. Scale bar: 1 μm.

We next explored if U-ExM could increase resolution in the region of the TAC close to the kDNA. We immunostained near non-fixed cells using a monoclonal antibody for TAC102 and anti-HA antibody for TAP110 and stained the DNA with DAPI. Confocal microscopy of the expanded cells showed that TAP110 is closer to the kDNA than TAC102 (Figure 5E).

### Importance of the TAC for TAP110 localization

In order to test if the kDNA is required for proper localization of TAP110 we created a dyskinetoplastic cell line that retains the TAC after loss of kDNA. This was achieved through depletion of p197 in ɣL262P bloodstream form cells, which are able to survive without kDNA [48]. P197 RNAi disrupts the TAC creating a population of dyskinetoplastic cells. We then can release p197 RNAi by removal of tetracycline from the medium and let the cells recover which, as previously shown, eventually leads to a reassembly of the TAC (simplified model Figure 6A) [36]. To monitor TAP110 it was PTP-tagged endogenously in the ɣL262P p197 RNAi cells. Five days post RNAi induction when 100% of the cells were dyskinetoplastic. In Immunofluorescence microscopy we observed that in about 55% of the cells TAC102 and TAP110 - while still colocalizing - seemed to mislocalize in the mitochondrion. Furthermore we observed a loss of TAC102 and TAP110 in about 40% of the cells, while 5% had a smaller signal for TAC102 and TAP110 (Figure 6B-C)). After re-expression of p197 (RNAi released) we detected that TAC102 and TAP110 signals returned to wild type localization and intensity (Figure 6B). Quantification of the experiment (Figure 6D-E) shows that TAC102 and TAP110 behave the same in the course of the recovery experiment.

**Figure 6:**
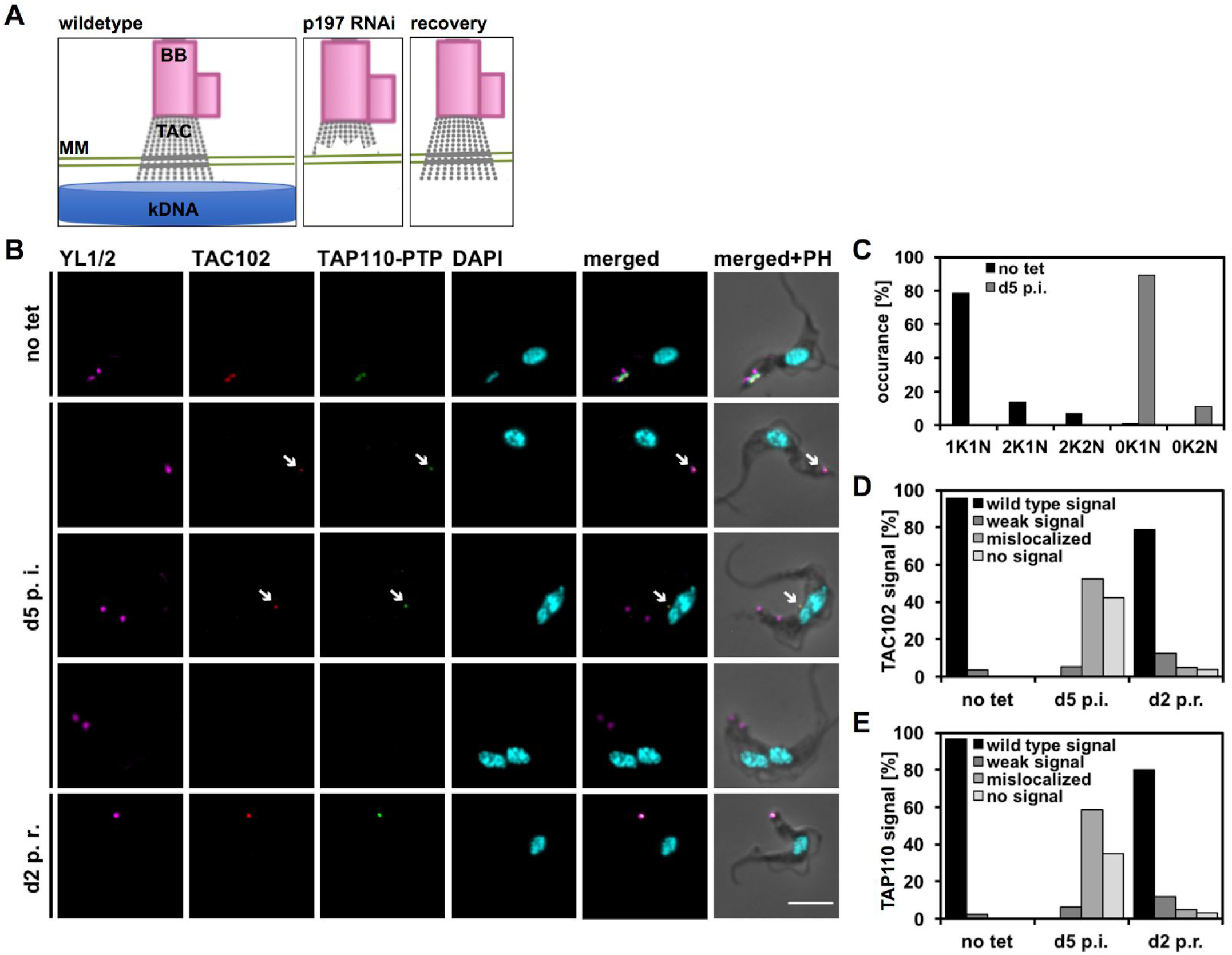
Recovery of TAP110 in ɣL262P p197 RNAi TAP110-PTP BSF cells. **A)** Depletion of p197 by RNAi in ɣL262P cells leads to loss of the TAC and kDNA; when RNAi against p197 is released in the same cells the TAC reassembles. **B)** Immunofluorescence microscopy of the ɣL262P p197 RNAi recovery experiment. Non-induced (no tet), induced cells at day five post induction (d5 p.i.) and cells at day two post recovery (d2 p.r.) are shown. Scale bar: 5 μm. Arrows point towards weak and mislocalized TAC102 and TAP110 signals. **C)** Quantification of the relative occurrence of kDNA networks and nuclei from experiment shown in B). A Minimum of n≥150 cells per time point were analyzed. **D)** Quantification of TAC102 signals from the experiment shown in B) (n≥150 for each time point). **E)** Quantification of TAP110 signals from the experiment shown in B) (n≥150 for each time point). BB, basal body; K, kDNA; N, nucleus, PH, phase contrast; TAC, tripartite attachment complex.

### TAP110 is insoluble in kDNA containing cells

The mitochondrial protein TAC102 and other TAC components can be partially solubilized by digitonin extractions as shown in earlier studies [30,34,36]. At a concentration of 0.025% digitonin cytoplasmic components can be separated from crude organellar structures containing the mitochondrial organelle. This crude mitochondrial pellet can then be lysed with 1% digitonin to solubilize mitochondrial proteins and partially also TAC proteins. We performed this extraction with the uninduced ɣL262P p197 RNAi cell line, which still contained the mitochondrial genome, and with a dyskinetoplastic version of this cell line. In the kDNA containing cells TAP110 was not soluble (Figure 7A), whereas in the dyskinetoplastic cells TAP110 could be partially solubilized (Figure 7B). We then performed blue native PAGE with extracts from wild type and dyskinetoplastic cells (Figure 7C). For TAC102 and another TAC protein TAC40, a component of the differentiated membranes, we detected protein complexes of around 440 kDa for TAC102 and between 500 and 750 kDa for TAC40. These sizes correspond to the complexes observed in earlier studies [36]. As TAP110 was not soluble in wild type cells, we only detected a protein complex for dyskinetoplasic cells. The protein complex for TAP110 was bigger than 669 kDa.

**Figure 7:**
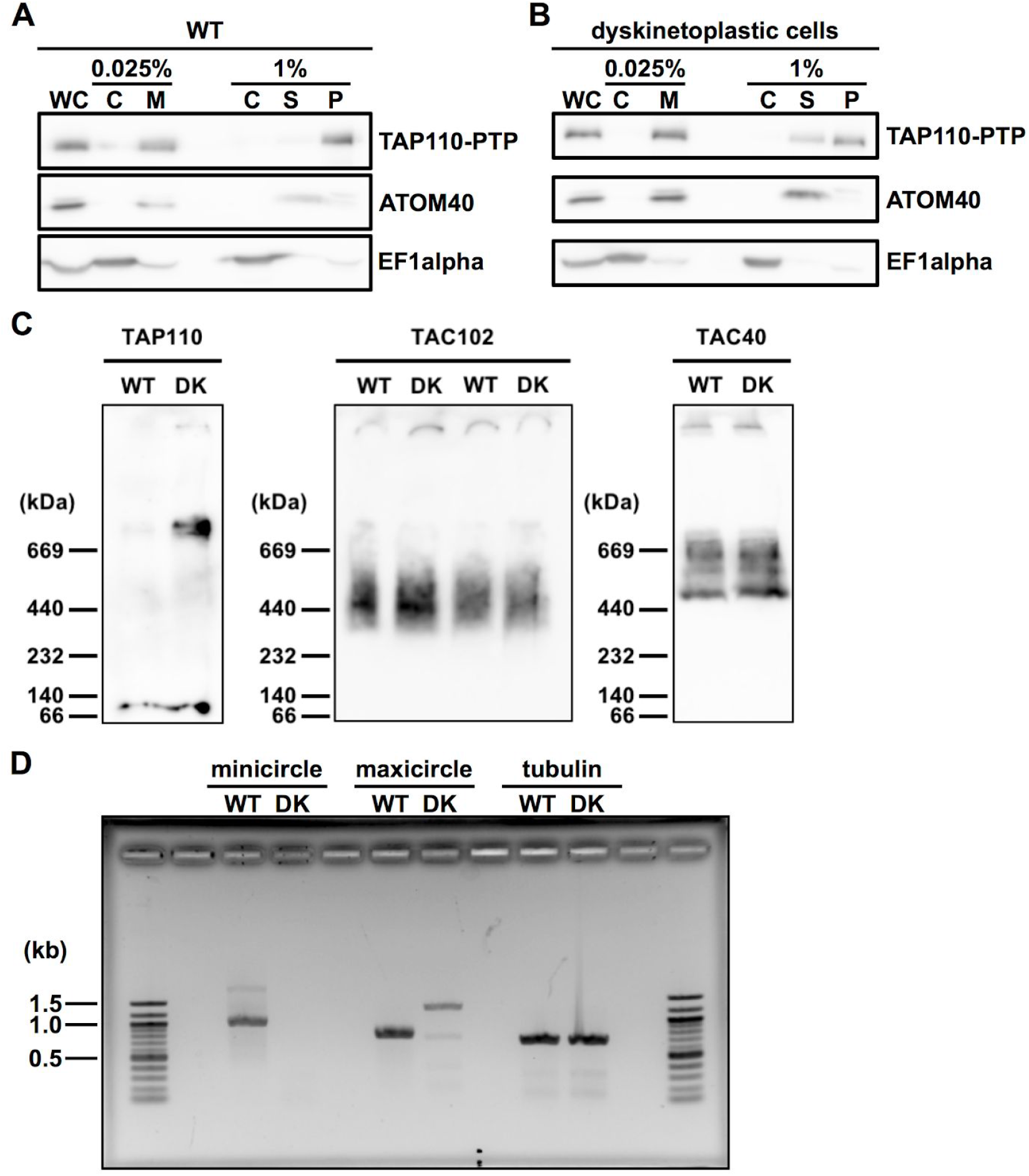
Biochemical analysis of TAP110 by western blot and blue native PAGE. **A)** Western blot of different digitonin extraction fractions obtained from wild type (WT) ɣL252P p197 RNAi TAP110-PTP BSF cells. ATOM40, mitochondrial marker; EF1alpha, cytosolic marker. **B)** Western blot of different digitonin extraction fractions obtained from dyskinetoplastic (DK) ɣL252P p197 RNAi TAP110-PTP BSF cells. **C)** Blue native PAGE from wild type and dyskinetoplastic cells lysed and soluble mitochondrial fractions. TAP110-PTP, TAC102 and EF1alpha were detected as described in Figure 2. TAC40-HA was detected with an anti-HA antibody and ATOM40 with an anti-ATOM40 antibody. **D)** PCR products using wild type (WT) or dyskinetoplastic cells (DK) gDNA and mini-, maxicircle or tubulin primers. C, cytosolic fraction; DK, dyskinetoplastic; M, mitochondrial fraction; P, insoluble mitochondrial fraction; S, soluble mitochondrial fraction; WC, whole cells; WT, wild type.

To verify dyskinetoplasticity of the cells, we performed PCRs on DNA extracted from wild type and dyskinetoplastic cells (Figure 7D). For this, we used primers designed to amplify whole minicircles and the gene p38 on the maxicircles. We didn’t detect any minicircle DNA in the dyskinetoplastic cells. With wild type DNA, we obtained the expected product of 767 bp in the maxicircle PCR, whereas for the dyskinetoplastic cells we obtained a product of around 1200 bp. We sequenced that PCR product and verified that it is not a product from amplification of maxicircle DNA but rather a product from amplification of an intergenic region of the nuclear genome. There was a very weak band of around 800 bp visible as well, however we were unable to validate this in a repetition of the same PCR (data not shown). Based on the PCR we believe that the dyskinetoplastic cells indeed are kDNA free, or at least minicircle free.

### TAP110 association with flagella

We used detergent extraction with Triton-X 100 to isolate flagella as described previously [28,30]. To obtain the flagellar fraction after lysis of the cells, the cytoskeletons were treated with a calcium chloride buffer that depolymerizes the subpellicular microtubules [49] and contained DNaseI in case of the +DNase samples. By centrifugation we separated the depolymerized subpellicular corset from the flagella. For comparison wild type and dyskinetoplastic cells were used for the extraction. The isolated flagella were used to perform immunofluorescence microscopy (Figure 8A-C). We stained for TAC102, TAP110 and the DNA as described above in Figure 1 and 2. In flagella obtained from wild type cells without DNaseI treatment, we observed that 59% had a signal for TAC102 and TAP110 and 26% of the flagella had a signal only for TAC102 (Figure 8A+B). Whereas in DNaseI treated flagella from wild type, only 24% had a signal for TAC102 and TAP110 and 63% had just a signal for TAC102 (Figure 8A+B). When we analyzed flagella obtained from dyskinetoplastic cells (without DNaseI treatment), we quantified that 90% showed a signal for TAC102 and TAP110 and 6% of the flagella had a signal for TAC102 only (Figure 8A+C). Although the dyskinetoplastic cells don’t contain kDNA, we performed an extraction including DNaseI treatment and observed that in this case 68% of the flagella had a signal for TAC102 and TAP110 and 19% of the flagella had only a signal for TAC102 (Figure 8A+C).

**Figure 8:**
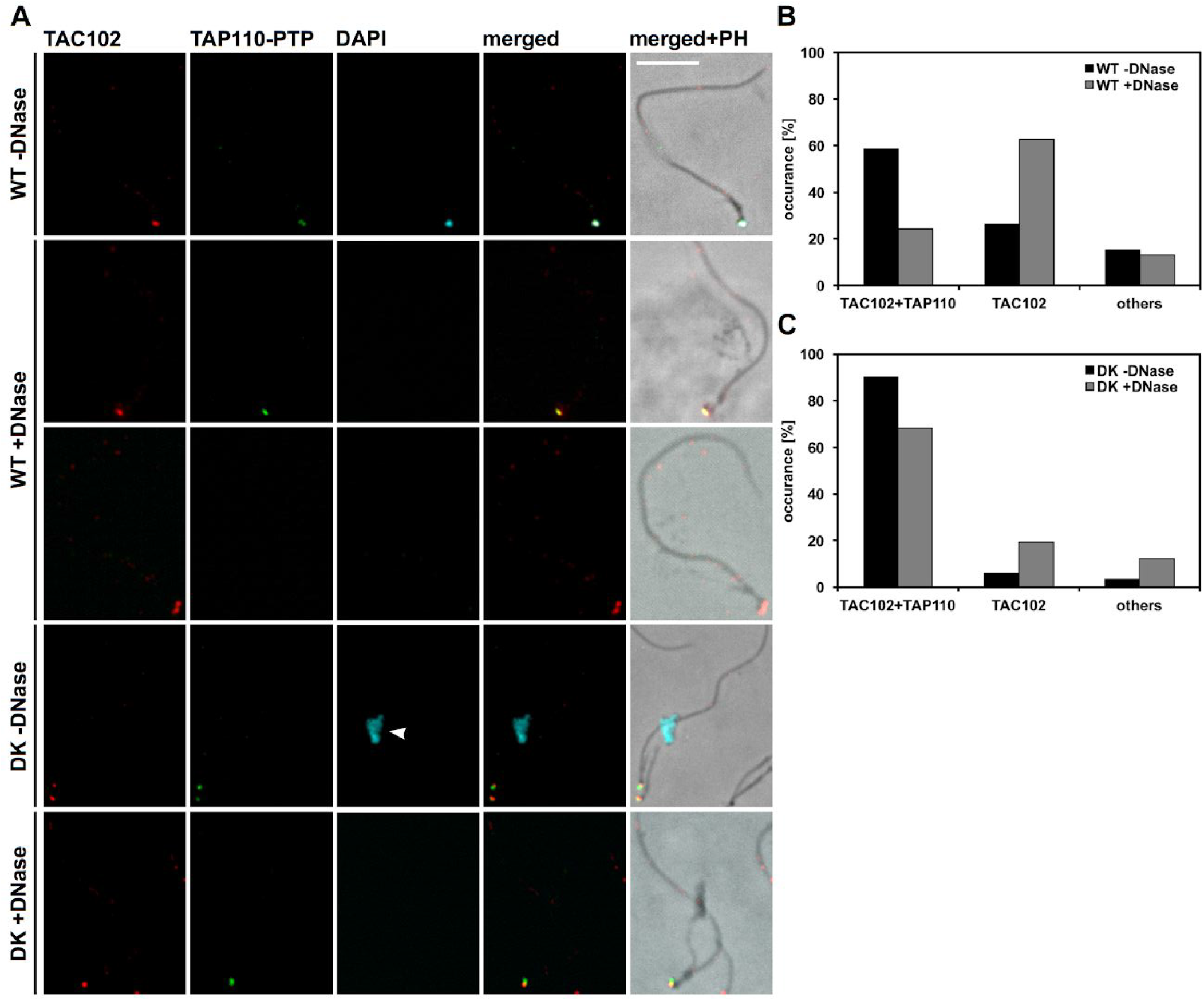
Localization of TAP110-PTP in flagellar extracts of wild type (WT) and dyskinetoplastic (DK)ɣL262P p197 RNAi TAP110-PTP BSF cells. **A)** Immunofluorescence microscopy images of extracted flagella either without DNaseI (-DNase) or with DNaseI treatment (+ DNase). Images were produced a s described in Figure 1. Scale bar: 5 μm. Arrowhead points towards nuclear DNA. **B)** Quantification wild type flagella of experiment shown in A). Flagella with TAC102 and TAP110-PTP signals (TAC102+TAP110), flagella with TAC102 signal only (TAC102) and other flagella (others) were counted. **C)** Quantification of dyskinetoplastic flagella of experiment shown in A). Quantification was performed as described in B).

## Discussion

Based on a comparative phylogenetic analysis TAC102 and TAP110 share a common evolutionary history (Figure S1D) and similarly to other TAC components TAP110 is not found in Perkinsela an endosymbiotic kinetoplast without basal body and flagellum. Interestingly, both proteins are also absent from the *Bodo saltans* genome, while all other TAC orthologues can be found in this free living kinetoplastid. This might suggest, that while most of the TAC machinery is conserved in all Kinetoplastea, the components in proximity to the DNA have adapted to the different kDNA conformations or replication mechanisms. We previously described a number of criteria for proteins of the TAC [29]. First, a TAC component is localized between the kDNA disc and the basal body of the flagellum. To determine the precise localization of TAP110 we used STED super resolution microscopy and found it to be co-localized with TAC102 in the unilateral filament region inside the mitochondrion. We further established ultrastructure expansion microscopy (U-ExM) for insect form trypanosomes and evaluated the expanded cells by confocal immunofluorescence microscopy. In general, the typical morphology of a trypomastigote cell was retained during the expansion process (Figure 5) and we were able to elucidate structures such as the subpellicular microtubule array or the microtubule quartet at the basal body as well as the ninefold symmetry of triplet microtubules of the barrel shaped pro-basal body (Figure S4E). The quasi isotropic expansion was also confirmed by the analysis of nucleus and basal body shape (Figure S4). However, quantitative analysis of the nuclear DNA, kDNA and the basal body revealed some differences in the expansion factor. DNA containing compartments did not expand as well as structural entities such as the basal body. Interestingly, *in situ* the kDNA only expanded in the horizontal plane of the disc, while the height of the disc remained largely unchanged. The kDNA is mainly composed of catenated minicircles that are relaxed and predicted to be oriented perpendicular to the horizontal plane of the disc [11–14]. The height of the kDNA disc is determined by the size of each minicircle (1kb) and reaches its maximum, when the minicircles are completely stretched out, which is apparently already the case in the unexpanded cells and therefore does not allow further expansion. The diameter of the kDNA disc and its expansion however, depend on the packaging of the minicircles in the horizontal plane. Tight packaging as predicted in the model would thus likely allow for relaxation and expansion as we detected in our experiments. After we evaluated the use of U-ExM for *T. brucei* we applied it to elucidate the localization of TAP110 relative to TAC102 and the kDNA. While in STED superresolution microscopy TAC102 and TAP110 colocalized with a correlation coefficient of 0.83, U-ExM revealed that TAP110 is the kDNA proximal protein, which is in good agreement with our current hierarchical model of TAC assembly and the TAC102/TAP110 RNAi data (Figure 2, 4). Aside from their localization between the basal body and the kDNA previously characterized TAC components also remain associated with flagella during their isolation irrespective of the presence or absence of kDNA. This is only partially true for TAP110. If we remove the kDNA from the flagella during extraction by DNAseI treatment, the number of flagella that contain detectable levels of TAC102 and TAP110 decreases by more than half suggesting that some fraction of TAP110 requires the kDNA or other kDNA connected proteins to be stably associated with the flagella/TAC. Interestingly, this requirement is decreased in flagella isolated from cells that do not contain any kDNA to begin with (dyskinetoplastic cells, Figure 8). In these cells TAP110 still co-localizes with TAC102 and remains with the TAC/flagellum throughout the extraction suggesting that TAP110, similar to TAC102 and the other known TAC components, does not require kDNA for its proper localization in the TAC region [36].

The association of TAP110 with the kDNA is also evident from its biochemical behaviour during solubilization of the TAC complex. While TAC102 becomes partially soluble irrespective of the presence or absence of kDNA, TAP110 is completely insoluble as long as kDNA is present (Figure 7). We have also tried to express TAP110 in order to perform DNA binding assays but have been unsuccessful in producing soluble protein. The third criterion for TAC proteins relates to their function in kDNA segregation. RNAi based depletion of any of the TAC components leads to a characteristic missegregation and eventually kDNA loss phenotype [29,30,36]. While TAP110 fulfills the first two criteria, it does not seem to be required for kDNA maintenance (Figure 2). Although the depletion and overexpression of TAP110 does lead to a detectable changes in kDNA content (Figure 3; S2; S3), it does not show any of the TAC typical kDNA segregation phenotypes. This might be explained by the presence of other proteins with redundant function or the incomplete depletion of TAP110. The lack of a kDNA maintenance related phenotype upon TAP110 depletion is also supported by the high throughput phenotyping study by Alsford and colleagues [50]. Interestingly, the overexpression of TAP110 leads to a kDNA segregation phenotype with a more than twofold increase in replicated but not segregated kinetoplasts. To test if and how TAP110 overexpression might influence other proteins we quantified the proteome during the overexpression of TAP110 (Figure 3). Among the list of proteins that change in expression level are six mitochondrial proteins of which three were shown to be kDNA associated. One of the kDNA associated proteins, Tb927.11.6660, was also identified in the initial biochemical assay where we screened for TAC102 interactors (Figure S1). Tb927.11.6660 localizes at the kDNA and the nucleus, which is consistent with its predicted mitochondrial as well as nuclear localization sequences (Figure S5). While the nuclear localization is predominantly observed during nuclear S-phase, the protein seems to be at the kDNA during the entire cell cycle. It is tempting to speculate that TAP110 and Tb927.11.6660 might provide a link for the coordinated replication/segregation of the mitochondrial and nuclear genome.

## Material and Methods

### *T. brucei* cell culture conditions

Procyclic form (PCF, 29-13) *T. brucei* cells were cultured in semi-defined medium-79 (SDM-79) supplemented with 10% FCS, 15 μg/ml geneticin and 25 μg/ml hygromycin at 27°C. Bloodstream form (BSF) T. brucei New York single marker (NYsm) cells [42] and γL262P BSF cells [48] were cultured at 37°C and 5% CO_2_ in Hirumi-modified Iscove’s medium 9 (HMI-9) supplemented with 10% fetal calf serum (FCS) containing 2.5 μg/ml geneticin for NYsm, for γL262P with 2.5 μg/ml geneticin and 0.5 μg/ml puromycin.

### Transfections of *T. brucei* cells

For transfections, we dissolved 10 μg of linearized plasmid or PCR product in 100 μl transfection buffer (90 mM Na-phosphate pH 7.3, 5 mM KCl, 0.15 mM CaCl_2_, 50 mM HEPES pH 7.3) [51]. For BSF and PCF transfection we used 4×10^7^ and 10^8^ mid-log phase cells, respectively. Cells were resuspended in 100 μl transfection buffer containing the DNA, transferred into Amaxa Nucleofector cuvettes and transfections were conducted in the Amaxa Nucleofector II using program Z-001 (panel V 1.2 kV, panel T 2.5 kV, panel R 186 Ohm, panel C 25 μF) for BSF and program X-014 for PCF.

Then, we recovered the transfected cells for 20 h in medium without antibiotics. After recovery we selected for correct integration of the construct with appropriate antibiotics (5 μg/ml blasticidin, 2.5 μg/ml, geneticin, 2.5 μg/ml, hygromycin, 2.5 μg/ml phleomycin or 0.5 μg/ml puromycin for BSF cells and for PCF cells 10 μg/ml blasticidin oror 1 μg/ml puromycin). Expression of the RNAi and overexpression constructs we induced by addition of tetracycline (tet) to a final concentration of 1 μg/ml for BSF and PCF cells.

### DNA constructs

TAP110-PTP was created by amplification of the TAP110 ORF (Tb927.11.7590) positions 2242 to 2922 from genomic NYsm DNA and cloned between the ApaI and EagI (NEB) sites of the pLEW100 based PTP tagging vector [41]. We linearized the resulting plasmid with XcmI (NEB) prior to transfection. TAP110 RNAi targeting the ORF (positions 2081 to 2629) was cloned into a tet inducible RNAi vector [43] in two steps by cloning with the restriction enzymes BamHI HF, HindIII HF, XbaI and XhoI (NEB) to generate the later hairpin loop dsRNA for RNAi. The final plasmid was linearized with NotI HF (NEB) prior to transfection. The ORF of TAP110 amplified and inserted without the stop codon by cloning with the restriction enzymes HindIII HF and XhoI (NEB) into a modified pLew100 vector for overexpression [42,52]. For the Tb927.11.6660-PTP construct the ORF positions 2397 to 2805 were amplified as described above and cloned between the ApaI and EagI sites. We used SnaBI (NEB) to linearize the plasmid prior to transfection.

### Immunofluorescence analysis

To analyze the localization of TAP110, TAC102, and basal body proteins, we used immunofluorescence analysis as described previously [53]. Primary and secondary antibodies were diluted as follows: polyclonal rabbit-anti-Protein A (Sigma) detecting the PTP epitope 1:2000, rat YL1/2 antibody detecting tyrosinated tubulin as present in the basal body [54] 1:100000, monoclonal mouse TAC102 antibody [30] 1:2000, rabbit-anti-HA (Sigma) 1:1000, Alexa Fluor® 488 Goat-anti-Rabbit IgG (H+L) (Invitrogen), Alexa Fluor® 594 Goat-anti-Mouse IgG (H+L) (Molecular probes), Alexa Fluor® 647 Goat-anti-Rat IgG (H+L) (Life technologies) all 1:1000. We acquired the images with the Leica DM5500 B microscope (Leica Microsystems) and the 100× oil immersion phase contrast objective. Then, we used the LAS X software (Leica Microsystems) and ImageJ to analyze the images.

### Super-resolution 2D Stimulated Emission Depletion microscopy

TAP110 PTP tagged BSF cells were used to analyze the TAP110 and TAC102 localization in more detail with Stimulated Emission Depletion (STED) microscopy. Cells were spread on Nr. 1.5 cover glasses (Marienfeld), fixed, permeabilized and mounted as described before [36]. Polyclonal rabbit-anti-Protein A antibody (Sigma) and monoclonal mouse-anti-TAC102 antibody were used as described above. The Alexa Fluor® 594 goat-anti-Rabbit IgG (H+L) (Invitrogen) and the Fluor® Atto647N Goat-anti-Mouse IgG (H+L) (Molecular probes) were used 1:500 in 4% BSA and incubated for one hour. To compare the localization of the two proteins we used the SP8 STED microscope (Leica, with a 100× oil immersion objective and the LAS X Leica software). Images were acquired as z-stacks with a z-step size of 120 nm and a X-Y resolution of 37.9 nm. For the TAP110-PTP and the TAC102 signal the 594 nm and 647 nm excitation laser and the 770 nm depletion laser were used. Due to non-availability of suitable depletion laser, the DAPI signal was acquired with confocal settings. We deconvoluted the images with the Huygens professional software.

### Calculation of Pearson correlation coefficient of TAP110-PTP and TAC102 signals obtained from 2D STED microscopy

To calculate the Pearson’s R value, we first selected the kDNA region as region of interest and then we applied the colocalization script of the ImageJ software according to the manufacturers user guide. This was performed for 30 different selected kDNA regions for the TAC102 and TAP110 signal.

### SDS-PAGE and western blotting

To analyze presence and/or abundance of a protein of interest in whole cell lysates and fractions from digitonin and flagellar extraction we used western blot analysis. Samples were prepared as described previously [53]. Approximately 5×10^6^ cells were loaded per lane and SDS PAGE and transferred to PVDF membrane. blocking and detection of proteins of interest (PTP and HA tagged proteins, EF1alpha and ATOM40) was performed as described previously [53]. Alpha tubulin was detected by the monoclonal mouse anti-alpha tubulin antibody (1:20’000, Sigma) and the rabbit anti-mouse-IgG conjugated to horseradish peroxidase (1:10’000, Dako).

### Mass spectrometry and data analysis

Protein lysates from induced (day 2) and non induced TAP110 overexpressing whole cells were separated on 10% NOVEX gradient SDS gel (Thermo Scientific) for 8 min at 180V in 1x MES buffer (Thermo Scientific). Proteins were fixated and stained with a Coomassie solution (0.25% Coomassie Blue G-250 [Biozym], 10% acetic acid, 43% ethanol). The gel lane was cut into slices, minced, and destained with a 50% ethanol/50 mM ammoniumbicarbonate pH 8.0 solution. Proteins were reduced in 10mM DTT for 1h at 56°C and then alkylated with 50 mM iodoacetamide for 45min at room temperature in the dark. Proteins were digested with trypsin (Sigma-Aldrich) overnight at 37°C. Peptides were extracted from the gel using twice with a mixture of acetonitrile (30%) and 50 mM ammoniumbicarbonate pH 8.0 solution and three times with pure acetonitrile, which was subsequently evaporated in a concentrator (Eppendorf) and loaded on an activated C18 material (Empore) StageTip [55].

For mass spectrometric analysis, peptides were separated on a 50 cm self-packed column (New Objective) with 75 μm inner diameter filled with ReproSil-Pur 120 C18-AQ (Dr. Maisch GmbH) mounted to an Easy-nLC 1200 (Thermo Fisher) and sprayed online into an Orbitrap Exploris 480 mass spectrometer (Thermo Fisher). We used a 103-min gradient from 3% to 40% acetonitrile with 0.1% formic acid at a flow of 250 nL/min. The mass spectrometer was operated with a top 20 MS/MS data-dependent acquisition scheme per MS full scan. Mass spectrometry raw data were searched using the Andromeda search engine [56] integrated into MaxQuant software suite 1.5.2.8 [57] using the TriTrypDB-46_TbruceiTREU927_AnnotatedProteins protein database (11,203 entries). For the analysis, carbamidomethylation at cysteine was set as fixed modification while methionine oxidation and protein N-acetylation were considered as variable modifications. Match between run option was activated.

### Bioinformatics analysis

Contaminants, reverse database hits, protein groups only identified by site, and protein groups with less than 2 peptides (at least one of them classified as unique) were removed by filtering from the MaxQuant proteinGroups file. Missing values were imputed by shifting a beta distribution obtained from the LFQ intensity values to the limit of quantitation. Further analysis and graphical representation was done in the R framework [58] incorporating ggplot2 package in-house R scripts [59].

The mass spectrometry proteomics data have been deposited to the ProteomeXchange Consortium via the PRIDE [60] partner repository with the dataset identifier PXD019665 (Reviewer account details: Username; Password Uitu7a0S).

### Ultrastructure expansion microscopy (U-ExM)

*T. brucei* cells were processed as indicated above for immunofluorescence. The protocol was adapted after [61]. After the last PBS wash, 150 μl containing two million cells were settled for 20 minutes at room temperature on poly-D-lysine functionalized coverslips (12 mm, Menzel-Glaser). Coverslips were transferred into 24 well plates filled with in a solution of 0.7% Formaldehyde (FA, 36.5-38%, Sigma) with 1% acrylamide (AA, 40%, Sigma) in PBS and incubated five hours at 37°C. Cells were then prepared for gelation by carefully putting coverslips (cells facing down to the gelling solution) into a 45μl drop of monomer solution (Sodium Acrylate (SA, 97-99%, Sigma) 10% (wt/wt) AA, 0.1% (wt/wt) N,N’-Methylenebisacrylamide (BIS, SIGMA) in PBS) supplemented with 0.5% APS and 0.5% tetramethylethylendiamine (TEMED) on parafilm in a pre-cooled humid chamber. Gelation proceeded for five minutes on ice, and then samples were incubated at 37°C in the dark for one hour. Coverslips with gels were then transferred into a six well plate filled with denaturation buffer (200 mM SDS, 200 mM NaCl, and 50 mM Tris in ultrapure water, pH 9) for 15 minutes at room temperature. Gels were then detached from the coverslips with tweezers and moved into a 1.5-ml Eppendorf centrifuge tube filled with denaturation buffer, and incubated at 95°C for one 90 minutes. After denaturation, gels were placed in beakers filled with deionized water for the first round of expansion. Water was exchanged at least twice every 30 minutes at room temperature, and then gels were incubated overnight in deionized water. Next day, gels were washed two times 30 minutes in PBS and subsequently incubated on a shaker (gentle) with primary antibody diluted in 2% PBS/BSA for 3 hours at 37 °C. Gels were then washed in PBST three times for 10 minutes while gently shaking and subsequently incubated with secondary antibody solution diluted in 2% PBS/ BSA for ~3 hours at 37°C. Gels were then washed in PBST three times for 10 minutes while gently shaking and finally placed in beakers filled with deionized water for expansion. Water was exchanged at least twice every 30 minutes before gels were incubated in deionized water overnight. Gel expanded between 3.61× and 3.86× according to SA purity.

The expansion factor was determined by comparing the ratio of expanded basal body, kDNA and nucleus to non-expanded basal body, kDNA and nucleus. For unexpanded kDNA and nucleus measurements N=22 cells from immunofluorescence imagery were analysed. For unexpanded basal body measurements transmission electron microscopy imagery was used (N=12). For measurements on the expanded cells N=22 cells were analyzed. We selected only nearly perfect side-view kDNA for measurement of kDNA length in each cell. The diameter of the nucleus was determined by measuring the widest diameter observed in each cell. The diameter of the basal body was determined by using the plot profile tools of Fiji to plot the Gaussian distribution, then the distance between the first and the last peak of intensity was measured.

### Digitonin fractionations

To analyze biochemical properties of TAP110 we performed digitonin fractionation as described before [53]. 5×10^6^ cell equivalents of each fraction was used for the western blot analysis. The samples were boiled for 5 min at 95°C in Laemmli buffer for SDS PAGE.

### Blue native analysis

To detect protein complexes, blue native PAGE analysis was performed as described before [36]. In brief, 10^8^ cell equivalents for each extract was used for the analysis. Crude mitochondrial fractions (obtained by extraction with 0.025% digitonin) were lysed with 1% digitonin and centrifuged for 15 minutes at 12000 rpm. The supernatant was loaded on a blue native gel, protein complexes were separated by electrophoresis, then the gel was soaked in SDS running buffer and the proteins were transferred onto a PVDF membrane by semi dry Western blotting.

### Flagellar extraction

For flagellar preparation [28], we performed the experiment as described before [53]. For DNase treatments, second extraction buffer was supplemented with DNaseI (Roche) to a final concentration of 100μg/ml.

### Phylogenetic analysis of TAP110 and TAC102

Phylogenetic analysis was done using the Phylo.fr package [62]. Sequences were aligned using the MUSCLE algorithm with standard settings [63]. Phylogenetic tree reconstruction was done using the PhylML 3.0 algorithm with standard settings [64]. Tree visualization and comparison was done using the Phylo.io tool [65].

### kDNA size measurements in DAPI stained cells

To measure the change of kDNA networks size upon depletion of TAP110, we spread approximately 10^6^ uninduced or cells at day three post induction onto slides. The experiment was performed as described previously [53]. In brief, the cells on a slide were fixed in cold methanol for 5 min at −20°C. Afterwards the slides were washed with PBS and then mounted with ProLong® Gold Antifade Mounting Medium containing DAPI. The images were acquired with a 100x oil immersion objective and analyzed using the ImageJ software. We measured the particle size in arbitrary units (a.u.).

## Acknowledgement

For the BBA4 and YL1/2 antibodies we would like to thank Keith Gull. For the anti-mtHsp70 antibody we would like to thank Luise Krauth-Siegel. For the RNAi and overexpression vector backbones we would like to thank André Schneider. We acknowledge Bernd Schimanski and Laura Pfeiffer for technical assistance and Christoph Wenger for the help with blue native PAGE. We would like to thank Anneliese Hoffmann, and Irina Bregy for discussions.

## Author Contribution

Torsten Ochsenreiter (TO) and Simona Amodeo (SA) designed the experiments. SA performed the experiments and analyses. TO and SA wrote the manuscript. Ana Kalichava (AK) designed, performed and analysed the U-ExM experiments. Paul Guichard (PG) provided reagents and helped design U-ExM experiments. Eloïse Bertiaux-Lequoy helped design and assisted in U-ExM experiments, provided U-ExM data. Albert Fradera-Sola (AF) designed, performed and analysed the mass spectrometry experiments. Falk Butter (FB) designed and analysed the mass spectrometry experiments.

---

**Figure S1:**
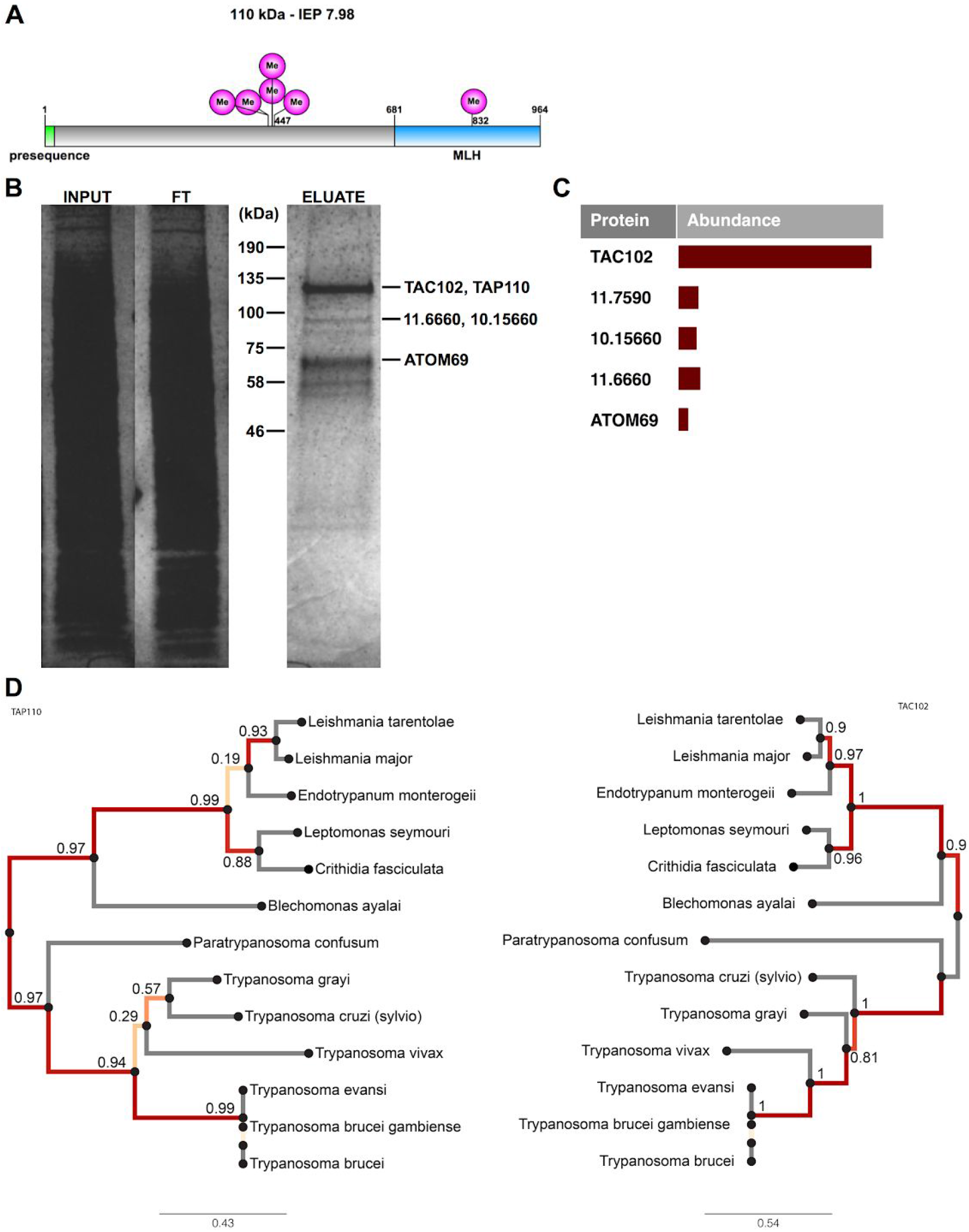
Protein domains, interaction partners and phylogeny of the TAC associated protein 110 (TAP110). **A)** Illustration of the TAP110 ORF. Highlighted in green, the mitochondrial-targeting sequence; in magenta, the posttranslational modification in form of methylarginine; in light blue, the similarity to the micronuclear linker histone protein of *Tetrahymena thermophila*. **B)** Coomassie gel of a PTP-TAC102 immunoprecipitation (IP) using IgG beads. FT, flow through. **C)** Abundance of TAC102 and its potential interaction partners detected in the eluate of the IP. **D)** A phylogenetic tree showing the conservation of TAC102 and TAP110 among Kinetoplastids. The scale bar indicates the number of amino acid substitutions.

**Figure S2:**
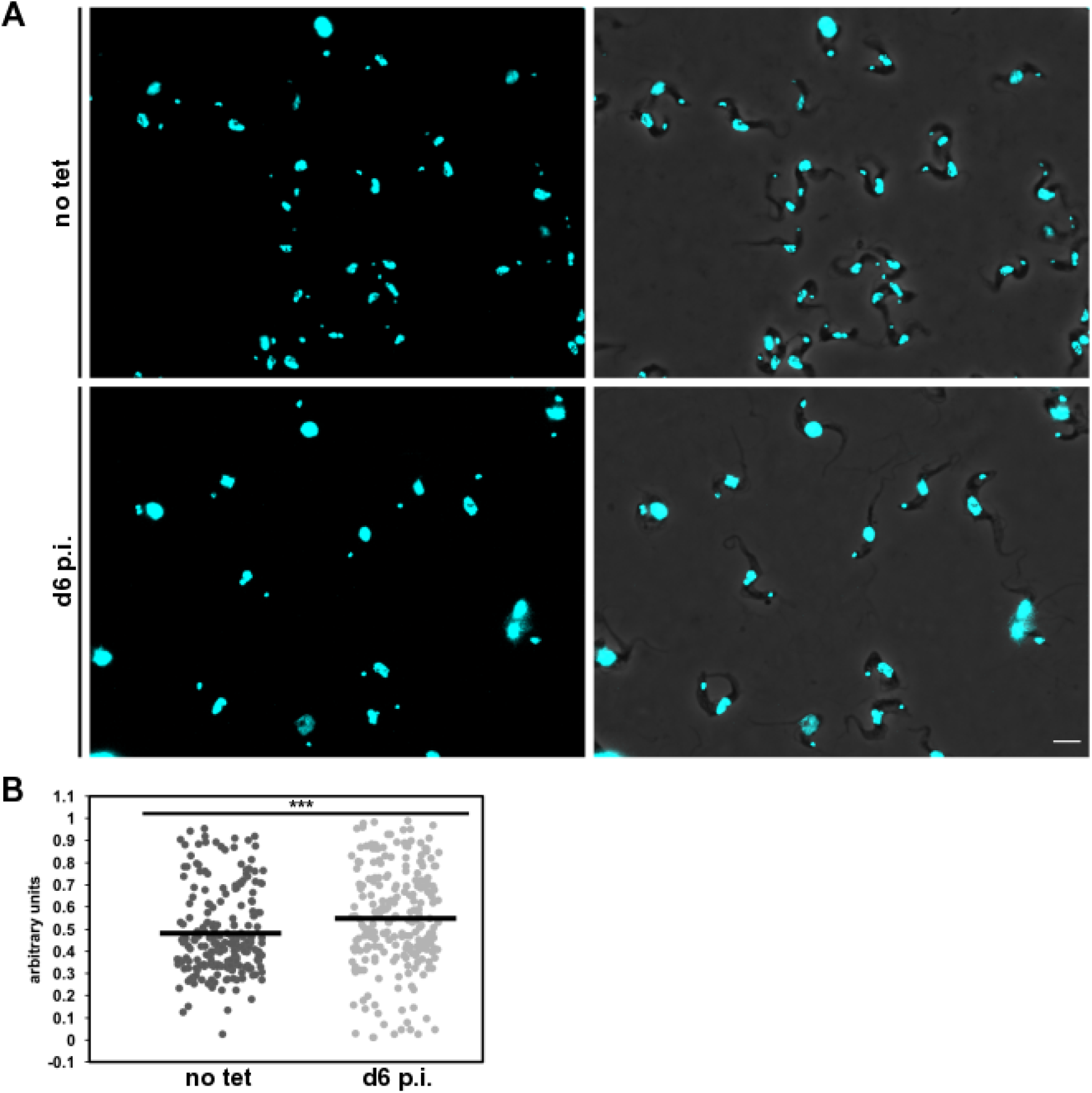
Measurement of kDNA content TAP110 RNAi in BSF cells. **A)** Immunofluorescence pictures from non-induced cells (no tet) and cells at day six of the TAP110 RNAi (d6 p.i.). Left side DAPI staining, right side overlay of DAPI with phase contrast. **B**) Size measurements (n ≥ 221 for each condition) of kDNA DAPI signals from uninduced (no tet; mean = 0.476 arbitrary units) and induced (d6 p.i.; mean = 0.547 arbitrary units) TAP110 RNAi BSF cells. Significance of difference in size was calculated by two-tailed unpaired t-test. *** = p ≤ 0.001.

**Figure S3:**
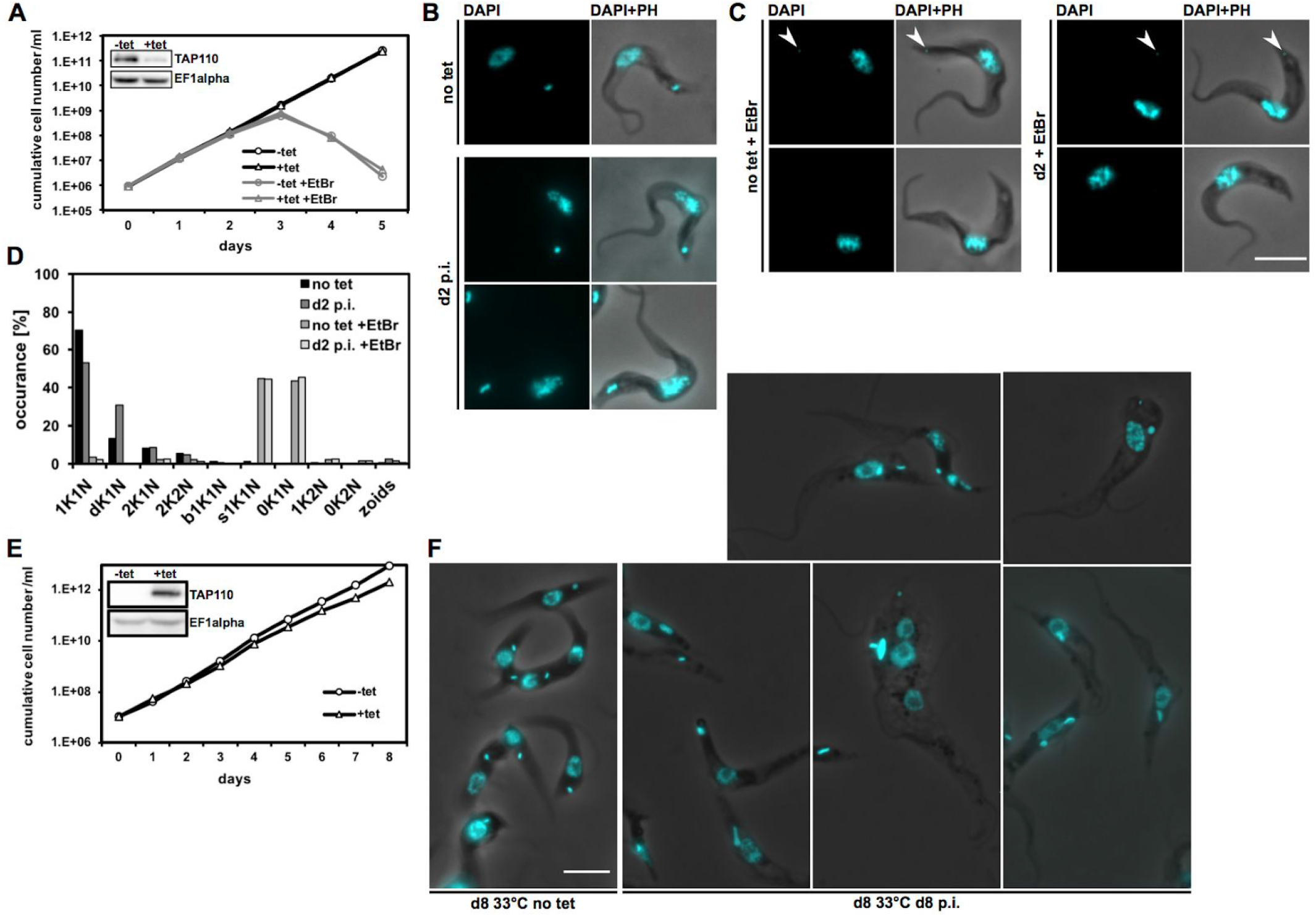
Ethidium bromide and heat stress experiment with the TAP110 RNAi BSF respectively TAP110 overexpression PCF cell line. **A)** Growth curve of tet inducible BSF TAP110 RNAi cells either grown without or with EtBr. The inset depicts a western blot, showing depletion of TAP110 at day one post induction. EF1alpha serves as a loading control. **B)** Representative fluorescence microscopy images of non-induced cells and cells at day two post induction which were not treated with EtBr. The nucleus and the kDNA were stained with DAPI. **C)** Representative fluorescence microscopy images of non-induced cells and cells at day two post induction which were treated with EtBr. Arrowheads point to small kDNA. **D)** Quantification of the relative occurrence of kDNA discs and nuclei. **E)** Growth curve of tet inducible TAP110 overexpression cells grown at 33°C. The inset depicts a western blot, showing expression of TAP110-HA at day one post induction. EF1alpha serves as a loading control. **F)** Representative fluorescence microscopy images of non-induced cells and cells at day eight post induction which were both grown for eight days at 33°C. Same staining procedure as in C was used. bK, big kDNA; dK, duplicating kDNA; K, kDNA; N, nucleus; PH, phase contrast; sK, small kDNA. Scale bar: 5 μm.

**Figure S4:**
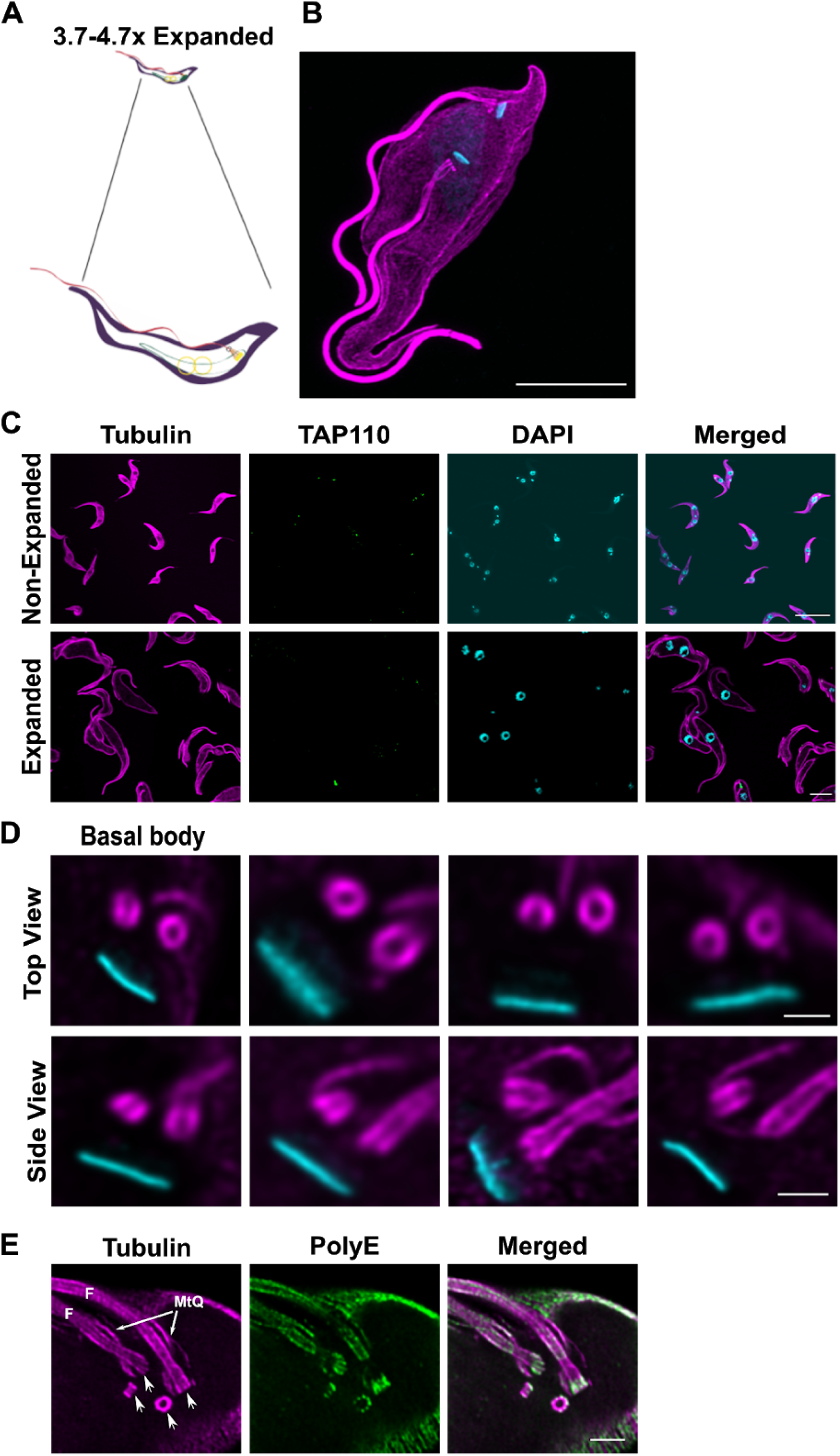
Isotropic expansion of *T. brucei* PCF whole cells and different subcellular structures by U-ExM. **A)** Schematic view of the expansion concept in *T. brucei*. **B)** Maximum intensity projection of an expanded PCF cell stained with anti-tubulin (magenta; Alexa Fluor 594) and DAPI (cayan) to illustrate a side view of a cell in a good orientation. Scale bar: 20 μm. **C)** Representative single plane images of expanded and non-expanded PCF cells stained with anti-tubulin (magenta; Alexa Fluor 594), anti-HA for TAP110 (green; Oregon Green 488) and DAPI for the kDNA and the nucleus (cayan). The cells were imaged by confocal microscopy and the imagery was deconvoluted by Huygens professional software. The z plane was selected based on nearly perfect view of the cells, therefore only one cell has TAP110 signal in the given single plane. Scale bar: 20 μm. **D)** Representative confocal images (deconvolved) of *T. brucei* basal bodies and kDNAs (stained as described above) from 2 different experiments. Scale bar: 2 μm. **E)** PCF cells stained for PolyE (green; Alexa Fluor 488) and anti-tubulin (magenta; Alexa Fluor 568) and imaged by confocal microscopy followed by HyVolution. Scale bar: 2 μm. Arrows point to the basal bodies. F, Flagella; MtQ, microtubule quartett

**Figure S5:**
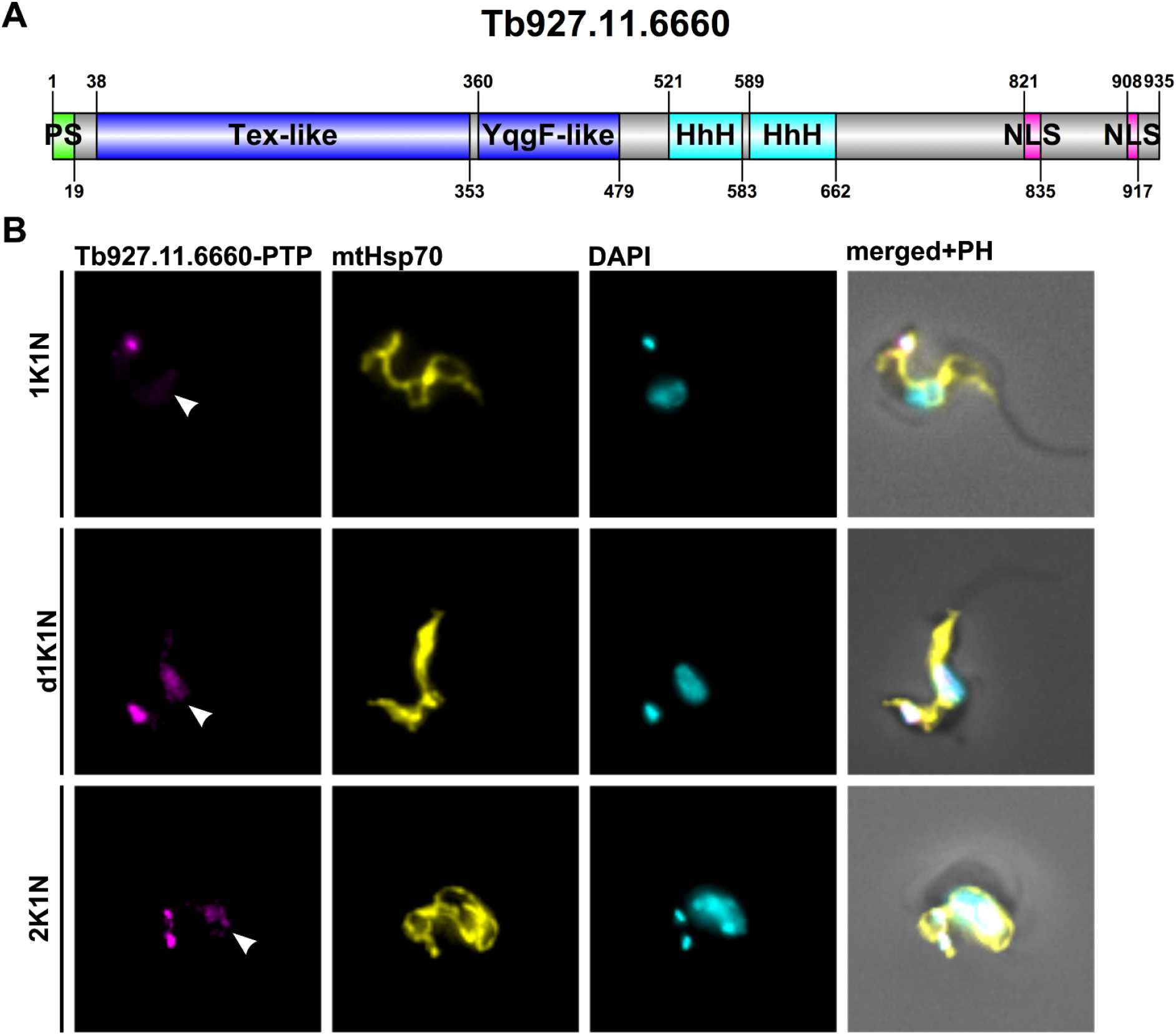
Protein domains and localization of Tb927.11.6660. **A)** Illustration of the Tb927.11.6660 ORF. Highlighted in green, the mitochondrial-targeting sequence; in dark blue, the Tex-like and YggF-like domains; in cyan, the helix-hairpin-helix motifs and in magenta, the nuclear localization signals. **B**) Immunofluorescence microscopy images of BSF cells expressing Tb927.11.6660-PTP during different stages of the cell cycle (1K1N, dK1N, 2K1N). Tb927.11.6660-PTP was detected with an anti-Protein A antibody. mtHsp70 with an anti-mtHsp70 antibody and the DNA with DAPI. K, kinetoplast; N, nulceus; PH, phase contrast.

